# The *LEPIS-HuR-TMOD4* axis regulates hepatic cholesterol homeostasis and accelerates atherosclerosis

**DOI:** 10.1101/2022.05.03.490400

**Authors:** Ping Lv, Hangyu Pan, Kexin Hu, Qinxian Li, Rongzhan Lin, Shaoyi Zheng, Zhigang Guo, Kai Guo

**Author notes:** Corresponding authors: Kai Guo, 1838 Guangzhou Avenue North, Guangzhou, Guangdong, People’s Republic of China), Zhigang Guo (1838 Guangzhou Avenue North, Guangzhou, Guangdong, People’s Republic of China), Shaoyi Zheng (1838 Guangzhou Avenue North, Guangzhou, Guangdong, People’s Republic of China).

## Abstract

Long noncoding RNAs (lncRNAs) play an important role in the entire progression of atherosclerosis. In this study, we identified an uncharacterized lncRNA, Liver Expressions by *PSRC1* Induce Specifically (*LEPIS*). The expression of *LEPIS* and its potential target tropomodulin 4 (*TMOD4*) in the liver of ApoE^-/-^ mice fed a high-fat diet was increased. An ApoE^-/-^ mouse model with the overexpression of *LEPIS* or *TMOD4* in liver was established, and we found that both *LEPIS* and *TMOD4* increased the burden of atherosclerosis and reduced hepatic cholesterol levels. Further study revealed that *LEPIS* and *TMOD4* affect the expression of genes related to hepatic cholesterol homeostasis, including proprotein convertase subtilisin/kexin type9 (*PCSK9*) and low-density lipoprotein receptor (*LDLR*), which are closely related to hypercholesterolemia. Mechanistically, human antigen R (HuR), an RNA-binding protein, was shown to be critical for the regulation of TMOD4 by LEPIS. Further, we found that overexpression of *LEPIS* promoted the shuttling of HuR from the nucleus to the cytoplasm, enhanced the stability of *TMOD4* mRNA, and in turn promoted the expression of TMOD4. In addition, TMOD4 was found to affect intracellular cholesterol levels through PCSK9. These results suggest that the *LEPIS-HuR-TMOD4* axis is a potential intervention target for hepatic cholesterol homeostasis and atherosclerosis.

**Graphical abstract:** 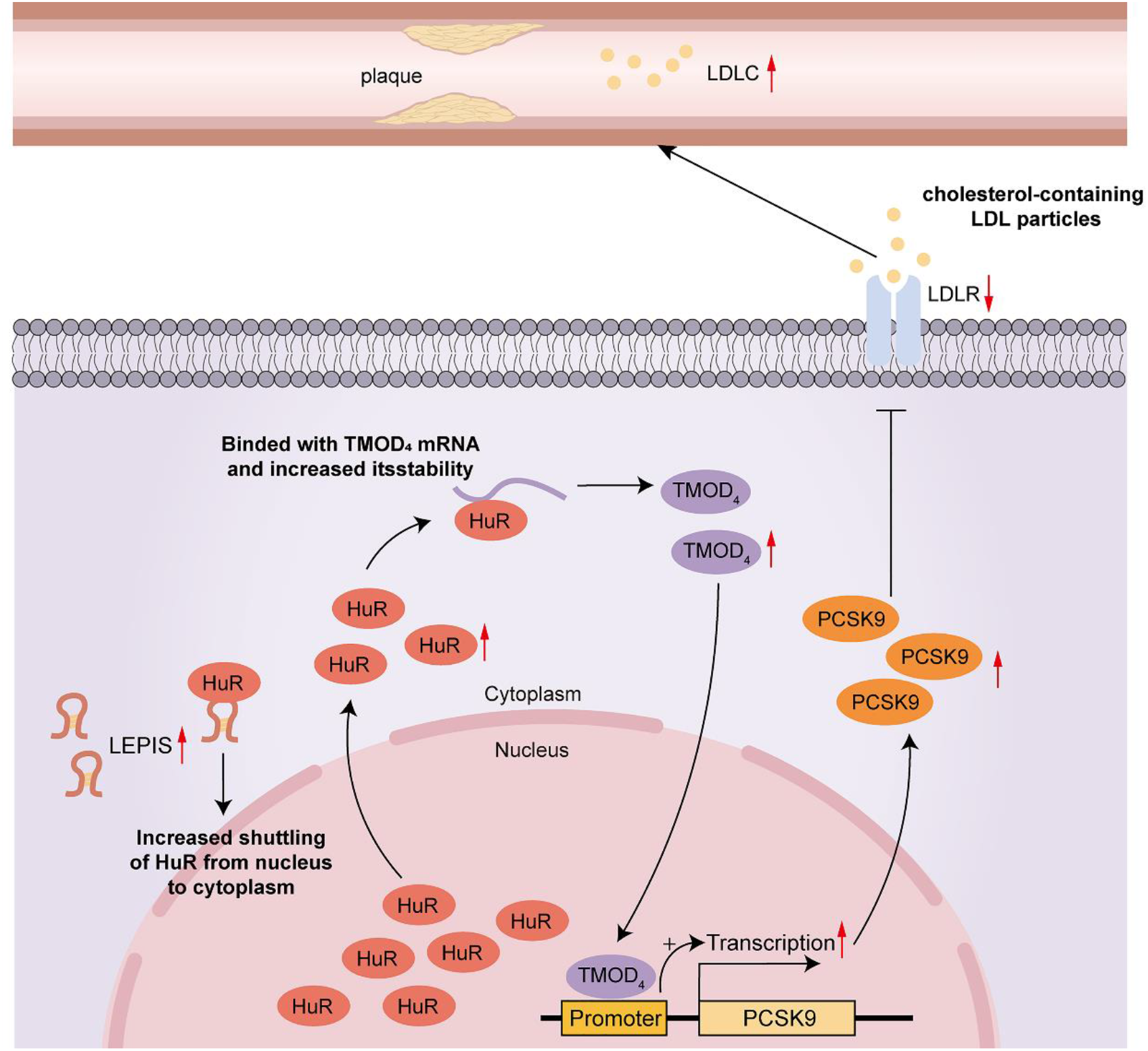

## INTRODUCTION

Cardiovascular disease is the greatest contributor to morbidity and mortality worldwide, especially within and across the WHO European Region(Townsend, Kazakiewicz et al., 2022). The underlying pathogenesis of atherosclerosis (AS) involves the disturbance of lipid metabolism and maladaptive immune responses to chronic inflammation of the arterial wall. Sites of laminar flow disturbance are prone to endothelial damage, and subendothelial accumulation of low-density lipoprotein can occur as a result. Oxidatively-modified lipids and low-density lipoproteins drive the infiltration of intimal immune cells, and lipid-laden macrophage-like foam cells form early atherosclerotic lesions(Weber & Noels, 2011). Multiple lines of evidence suggest that the degree and duration of exposure to LDL determines the risk of atherosclerotic vascular disease and its complications. Therefore, maximally lowering the level of LDL-C in plasma is still the essential step for preventing and treating ASCVD.

A large meta-analysis of 26 random clinical trials suggested that more intensively lowering LDL-C would provide additional protection for high-risk patients(Baigent, Blackwell et al., 2010). The efficacy of PCSK9 inhibitors in lowering LDL is only 50-60%(Tokgozoglu & Libby, 2022). Studies have confirmed that there is a residual risk of ASCVD for individuals treated with sufficient lipid-lowering therapies, such as lifestyle improvements, statin therapy, and PCSK9 inhibition(Hussain & Ballantyne, 2021, Wang, Li et al., 2022). In addition, individuals are pursuing lipid-lowering methods that exhibit better lipid-lowering effects and can be used less frequently than current methods(Tokgozoglu & Libby, 2022). A new era of targeted lipid-lowering therapy is gradually coming, and this also provides the possibility of individualized treatment for patients with different risk levels(Tokgozoglu & Libby, 2022). Therefore, it is essential to seek new ways to reduce AS-related mortality and morbidity.

Less than 2% of transcripts have the capacity to encode proteins in the human genome, and there are many noncoding RNAs in the human genome. Long noncoding RNAs (lncRNAs) are a class of noncoding RNA molecules (ncRNAs) comprising more than 200 nucleotides, which are highly tissue-specific and usually not very conserved. Emerging evidence indicates that lncRNAs can modulate gene expression via diverse mechanisms and contribute to the development of various diseases. Some lncRNAs have been demonstrated to work as regulators of the key steps in the progression of atherosclerosis, and these steps include cholesterol metabolism, the inflammation cascade, and apoptosis(Meng, Pu et al., 2020). However, there are still many lncRNAs that urgently need to be studied. Our previous study showed that PSRC1, which is an atheroprotective gene, could significantly induce the downregulation of a particular lncRNA expression in liver, and this is called Liver Expressions by PSRC1 Induce Specifically (*LEPIS*)(Guo, Kai et al., 2018, Mengqiu, Wei et al., 2020). However, the pathobiological roles of *LEPIS* have so far remained unknown and warrant investigation. The purpose of this study was to investigate the mechanism by which *LEPIS* regulates cholesterol metabolism and the progression of AS and to provide new evidence for effective strategies that further lower the levels of lipids in plasma or prevent and treat AS.

In our study, the role of *LEPIS* and its pathways were thoroughly investigated. We found that ① the overexpression of *LEPIS* in the liver promotes the occurrence of AS and reduces plaque stability; ② *LEPIS* interacts with the RNA-binding protein (RBP) HuR to regulate the expression of *TMOD4*; ③ *LEPIS* promotes the shuttling of HuR from the nucleus to the cytoplasm, and HuR binds to *TMOD4* mRNA and regulates its stability, which is a classic way that HuR exerts biological effects; and ④ we also identified the binding region of HuR to LEPIS and the binding region of TMOD4 to the *PCSK9* promoter.

## RESULTS

### *LEPIS* and its potential target *TMOD4* are involved in cholesterol homeostasis and atherosclerosis

The expression of *LEPIS* in the liver of ApoE^-/-^ mice fed a chow diet and a high-fat diet (HFD) for 12 weeks was detected, and the expression of *LEPIS* in the liver of ApoE^-/-^ mice fed a HFD was significantly upregulated compared with that observed in ApoE^-/-^ mice fed a chow diet (Figure 1A). Then, a series of bioinformatics analyses found that *LEPIS* is approximately 956 bp in length and has 6 exons. *LEPIS* exhibits a weak ability of encoding proteins, and it is well conserved among different species. In addition, *LEPIS* has the ability of binding DNA, RNA and proteins **(Supplementary Figure 1A-D)**. Furthermore, knockdown of *LEPIS* using small interfering RNA (siRNA) resulted in a marked increase in cholesterol levels in HepG2 cells (Figure 1B). The above research and bioinformatics results indicate that *LEPIS* is related to the cholesterol homeostasis of hepatocytes and is possible to affect the occurrence and development of AS by regulating cholesterol metabolism in the liver.

**Figure 1.**
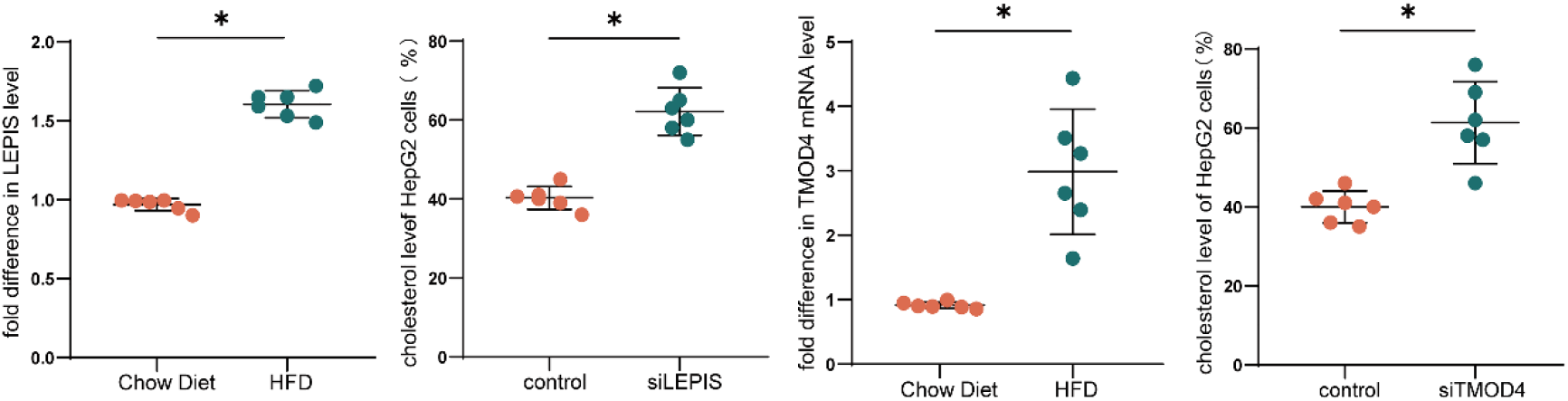
*LEPIS* and its potential target *TMOD4* are closely related to cholesterol metabolism and AS. (A) Quantitative real-time PCR analysis of *LEPIS* expression in the livers of ApoE^-/-^ mice fed a chow diet (n=6) or a HFD (n=6); (B) Intracellular cholesterol levels in HepG2 cells after silencing *LEPIS*; (C) Quantitative real-time PCR analysis of *TMOD4* expression in the livers of the ApoE^-/-^ mice fed a chow diet (n=6) or a HFD (n=6); (D) Intracellular cholesterol levels in HepG2 cells after silencing *LEPIS* (*P<0.05).

Studies have shown that lncRNAs can exert their biological effects by regulating the expression of nearby genes in both cis and trans conformations(Meng et al., 2020). Therefore, we analyzed the potential downstream targets of *LEPIS*, and the results showed that *LEPIS* and *TMOD4* are located on the same chromosome, are close to each other and have fragments that overlap with *TMOD4* **(Supplementary Figure 1E).** Thus, *LEPIS* may function by affecting the expression of *TMOD4*. After mice were fed with a chow diet or HFD for 12 weeks, RT–qPCR and Western blot were used to detect the expression of *TMOD4* in the livers of ApoE^-/-^ mice, which showed that the expression of *TMOD4* in the ApoE^-/-^ mice fed a HFD was significantly increased (Figure 1C). After knockdown of *TMOD4* expression was performed in HepG2 cells with siRNA, the cholesterol levels of the cells were significantly increased (Figure 1D). In conclusion, *LEPIS* may play a role in regulating the cholesterol metabolism of livers and affecting the occurrence and development of atherosclerosis by regulating the expression of *TMOD4*.

### Overexpression of *LEPIS* and *TMOD4* in the liver promotes AS

After AAV was injected in the tail vein of mice and the HFD was continued for 4 weeks, the total RNA in the liver tissues of mice in the three groups was extracted and determined. The expression of *LEPIS* and *TMOD4* in the livers of AAV-LEPIS mice was upregulated significantly, and the expression of *TMOD4* in the livers of AAV-TMOD4 mice was also upregulated, which indicated that the ApoE^-/-^ mouse model with *LEPIS* or *TMOD4* overexpression in the liver was successfully constructed (Figure 2A). In addition, we found that the expression of *LEPIS* in livers was equivalent in AAV-NC mice and AAV-TMOD4 mice (Figure 2A). In summary, an animal model with *LEPIS* or *TMOD4* overexpression in livers was successfully established. The baseline results are shown in **Supplementary Figure 2**.

**Figure 2.**
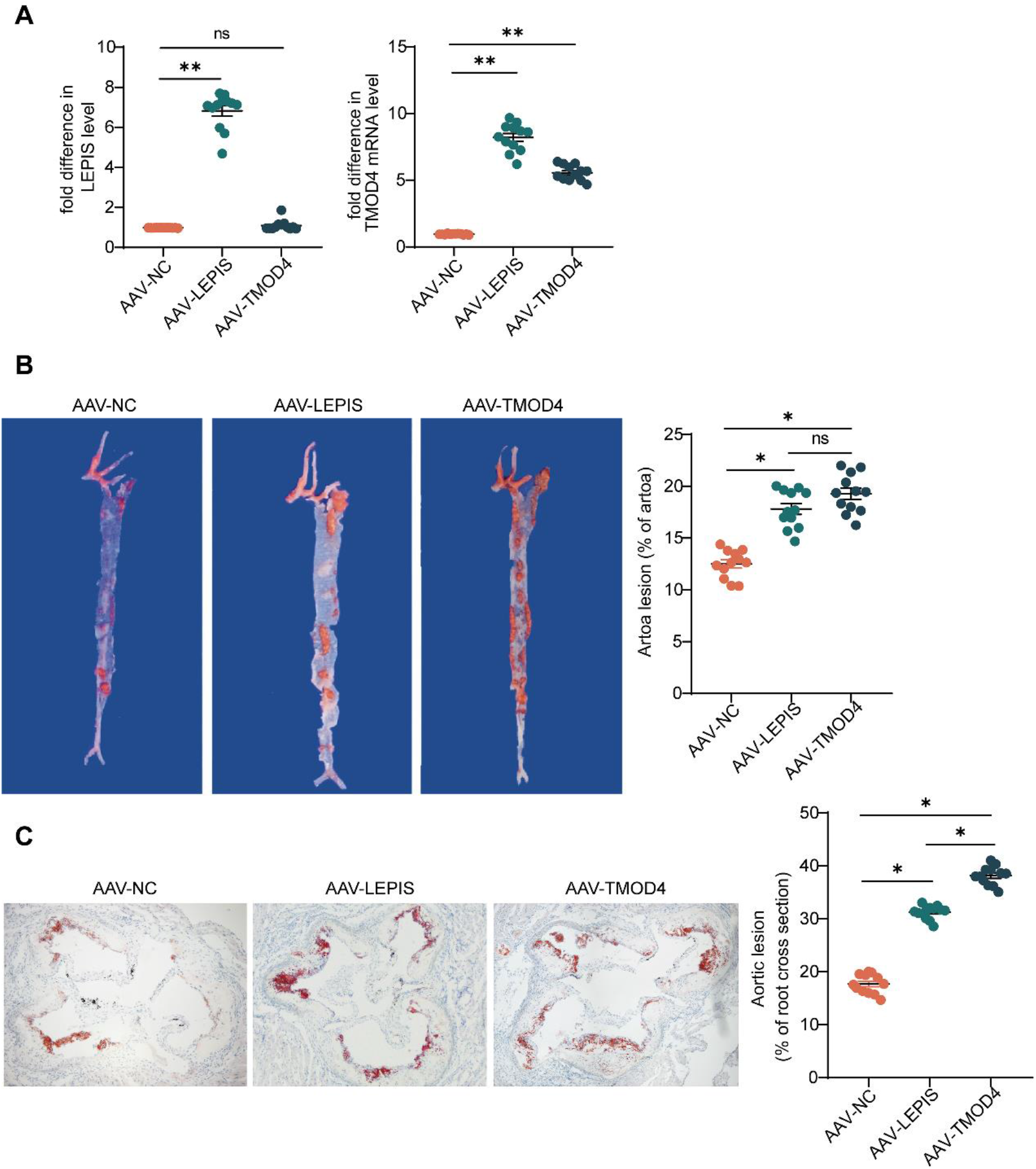
*LEPIS* and *TMOD4* promote AS. (A) Quantitative real-time PCR analysis of *LEPIS* expression (n=12) and *TMOD4* expression (n=12) in the livers of ApoE^-/-^ mice 4 weeks after tail vein injections of adeno-associated virus; (B) Oil red O staining of whole aorta from ApoE^-/-^ mice 4 weeks after tail vein injections of adeno-associated virus (n=12 in each group); (C) Oil red O staining of the cross-sections of the aorta from ApoE^-/-^ mice 4 weeks after tail vein injections of lentivirus (n=12 in each group, 100×).

The aortas of the above mice in the three groups were stained with oil red O to observe the burden of AS. It was suggested that compared with the AAV-NC mice, the ratio of aortic lesion/aorta in the AAV-LEPIS mice and AAV-TMOD4 mice were increased by 5.3% and 6.78%, respectively (Figure 2A). The ratio of aortic lesion/cross section of the aorta was increased by 13.53% and 20.43%, respectively (Figure 2B). These findings suggested that overexpression of *LEPIS* or *TMOD4* in the liver accelerates AS. HE staining, Masson staining, immunofluorescence staining and EVG staining were performed with the aortic roots from the above three groups of mice. Compared with AAV-NC mice, the cross section of the aorta in AAV-LEPIS mice and AAV-TMOD4 mice had a larger lipid core area, increased expression of CD68, decreased expression of a-SMA, and decreased elastic fibers (Figure 3). This suggested that increased expression of either *LEPIS* or *TMOD4* in the liver reduces plaque stability.

**Figure 3.**
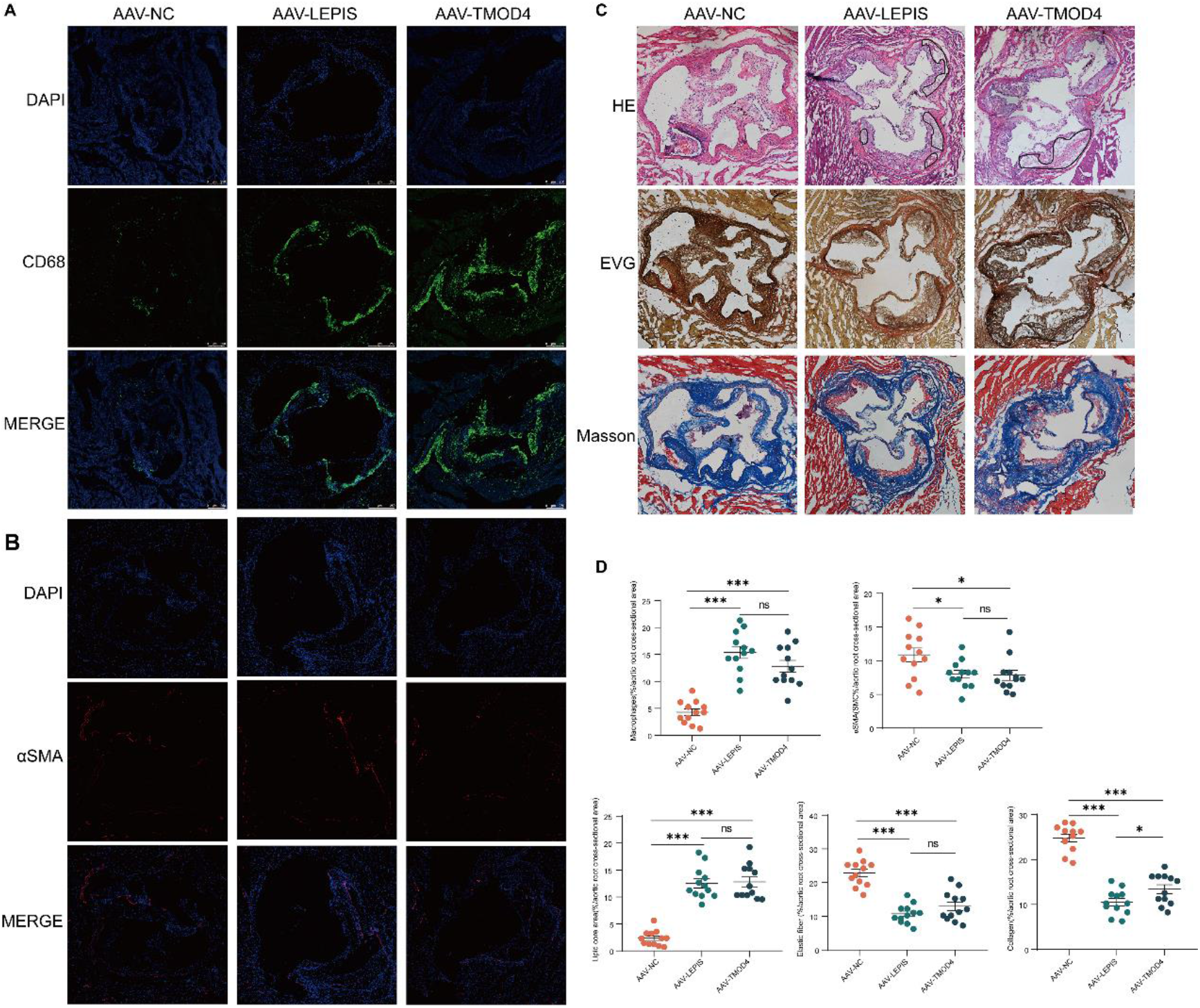
LEPIS and TMOD4 reduce the plaque stability. (A) Immunofluorescence for CD68 in the crossing sections of the aorta from ApoE-/- mice 4 weeks after tail vein injections of lentivirus (n=12 in each group, 50×); (B) Immunofluorescence for αSMA in the cross-sections of the aorta from ApoE-/- mice 4 weeks after tail vein injections of lentivirus (n=12 in each group, 50×); (C) HE staining, EVG staining, Masson staining of the crossing-sections of the aorta from ApoE-/- mice 4 weeks after tail vein injections of lentivirus (n=12 in each group, 100×); (D) Statistics of the above staining.

### *LEPIS* and *TMOD4* regulate the homeostasis of hepatic cholesterol

After 12 weeks fed with HFD, there were no significant differences in the body weight, liver weight, or liver weight/body weight among the three groups of mice **(Supplementary Figure 3)**. The peripheral blood serum of mice in the AAV-NC group, AAV-LEPIS group and AAV-TMOD4 group was collected, and the lipid levels were detected. The results showed that compared with AAV-NC mice, the levels of triglycerides (TGs) in AAV-LEPIS mice and AAV-TMOD4 mice were increased by 68.01% and 78.94%, respectively. The levels of total cholesterol (T-CHO) in AAV-LEPIS mice and AAV-TMOD4 mice were increased by 93.42% and 103.04%, respectively. The levels of high-density lipoprotein cholesterol (HDL-C) in AAV-LEPIS mice and AAV-TMOD4 mice were increased by 35.55% and 46.35%, respectively. The levels of low-density lipoprotein cholesterol (LDL-C) in AAV-LEPIS mice and AAV-TMOD4 mice were increased by 126.24% and 140.83%, respectively, and the above differences were significant (Figure 4A). It can be concluded that *LEPIS* or *TMOD4* overexpression in the liver increased the lipid levels in plasma.

**Figure 4.**
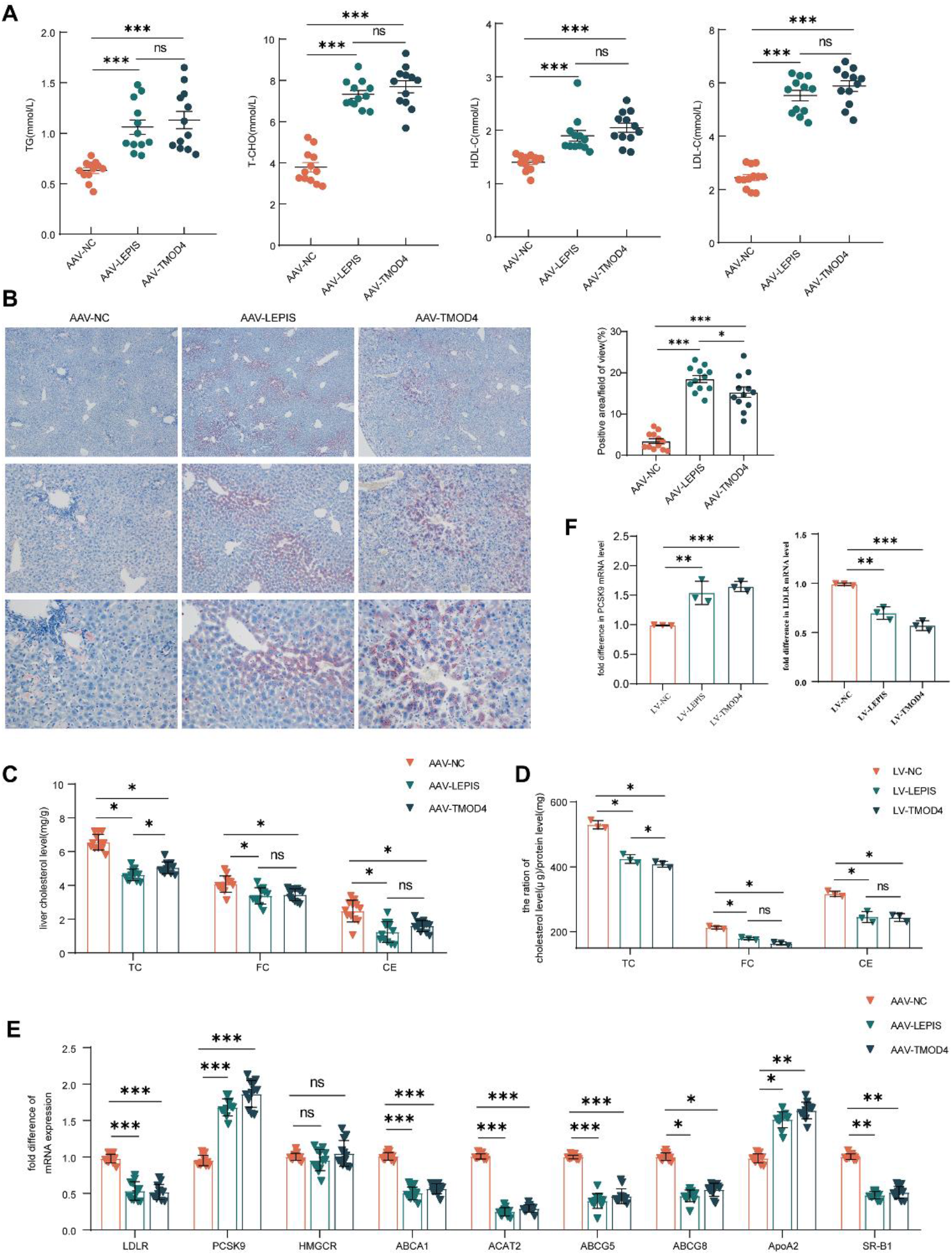
*LEPIS* and *TMOD4* regulate hepatic cholesterol homeostasis. (A) The levels of TG, TCHO, HDL-C and LDL-C in the plasma of ApoE^-/-^ mice in each group (n=12 in each group); (B) Oil red O staining of the livers of ApoE^-/-^ mice in each group (n=12 in each group, 40×, 100×, 200×); (C) Levels of TC, FC and CE in the livers of ApoE^-/-^ mice in each group (n=12 in each group); (D) Levels of TC, FC, and CE in HepG2 cells; (E) Expression of cholesterol metabolism-related genes in the livers of ApoE^-/-^ mice in each group (n=12 in each group);(F) *PCSK9* and *LDLR* expression in HepG2 cells after *LEPIS* or *TMOD4* upregulation.

The liver tissues from AAV-NC mice, AAV-LEPIS mice and AAV-TMOD4 mice were taken and stained with Oil Red O. The results showed that the triglycerides in the liver tissue of AAV-LEPIS mice and AAV-TMOD4 mice were significantly higher than those in AAV-NC mice (Figure 4B). At the same time, cholesterol levels in liver tissue were detected. The levels of total cholesterol (TC), free cholesterol (FC) and cholesterol esters (CE) in the livers of AAV-LEPIS mice and AAV-TMOD4 mice were lower than those of AAV-NC mice (Figure 4C). In vitro, LEPIS-overexpressing (LV-LEPIS) HepG2 cells and TMOD4-overexpressing (LV-TMOD4) HepG2 cells were constructed by transfection of lentivirus, and the multiplicity of infection (MOI) was 50 pfu/cells and 25 50 pfu/cells, respectively **(Supplementary Figure 4)**. Consistent with the in vivo experiments, the cholesterol levels of LV-LEPIS and LV-TMOD4 HepG2 cells were lower than those of LV-NC HepG2 cells (Figure 4D). This confirmed that *LEPIS* and *TMOD4* regulated the homeostasis of cholesterol in hepatocytes.

Cholesterol metabolism is mainly divided into the following aspects: cholesterol synthesis, cholesterol uptake, cholesterol efflux, and cholesterol esterification(Luo, Yang et al., 2020). To further elucidate how the overexpression of hepatic *LEPIS* and *TMOD4* regulates cholesterol metabolism, we examined the expression of representative genes that are related to hepatic cholesterol metabolism. First, we searched the *GeneCards* database for genes or molecules that were related to hypercholesterolemia and screened the top 10 genes for verification by RT-qPCR **(Supplementary Table 1)**. Apart from *HMGCR*, which is related to cholesterol synthesis, we found that the expression of other genes in AAV-LEPIS and AAV-TMOD4 mice were significantly different from that in AAV-NC mice. Specifically, the levels of *LDLR* in AAV-LEPIS mice and AAV-TMOD4 mice were downregulated, and *PCSK9* was upregulated. The levels of *ABCA1* were downregulated, the levels of *ApoA2* were upregulated, the levels of *ABCG5* were downregulated, and the levels of *ABCG8* were downregulated (Figure 4E). Considering the important roles of *PCSK9* and *LDLR* in cholesterol metabolism, we next validated the changes in expression of *PCSK9* and *LDLR* in LV-LEPIS and LV-TMOD4 HepG2 cells. The results were consistent with those of the in vivo experiments (Figure 4F). Thus, *LEPIS* and *TMOD4* affect the homeostasis of hepatic cholesterol by regulating genes that are related to cholesterol metabolism.

### *LEPIS* upregulates *TMOD4* and thereby increases the expression of *PCSK9* in hepatocytes

Reports that PCSK9 and LDLR are related to LDL-C levels in peripheral blood have emerged, and LDL-C is significantly reduced by PCSK9 inhibitors, which is an effective method for the clinical treatment of hypercholesterolemia and AS(Johns, Almonte et al., 2021, Karagiannis, Liu et al., 2018, Knowles, Howard et al., 2017, Tang, Li et al., 2020). To further investigate how *LEPIS* and

### *TMOD4* regulate cholesterol homeostasis in hepatocytes, we focused on how *LEPIS* and *TMOD4* affect the levels of PCSK9

There are many reports that lncRNAs exert their biological effects by regulating the transcription of nearby protein-coding genes(Meng et al., 2020). Considering that the *TMOD4* and *LEPIS* are both located on chromosome 1 with a portion of *LEPIS* overlapping with the 5′ portion of *TMOD4* **(Supplementary Figure 1)** and that *LEPIS* and *TMOD4* have the same effect on cholesterol homeostasis and AS, we further verified whether *LEPIS* regulates the expression of *TMOD4*. Consistent with the in vivo experiments (Figure 5A), overexpression of *LEPIS* with lentivirus in HepG2 cells resulted in significantly elevated mRNA and protein levels of TMOD4 (Figure 5B-C). In addition, the expression of PCSK9 was significantly increased, and the expression of LDLR was significantly decreased (Figure 5A-D). Furthermore, based on the overexpression of *LEPIS*, HepG2 cells (LV-LEPIS/si-TMOD4) with *LEPIS* overexpression and *TMOD4* knockdown were established. Compared with that in LV-LEPIS cells, the intracellular cholesterol levels were upregulated (Figure 5E), the expression of PSCK9 was downregulated, and the expression of LDLR was upregulated in LV-LEPIS/si-TMOD4 cells (Figure 5B-D). In conclusion, *LEPIS* regulates the expression of *PCSK9* in hepatocytes by targeting *TMOD4*, and this plays a role in regulating cholesterol metabolism in hepatocytes

**Figure 5.**
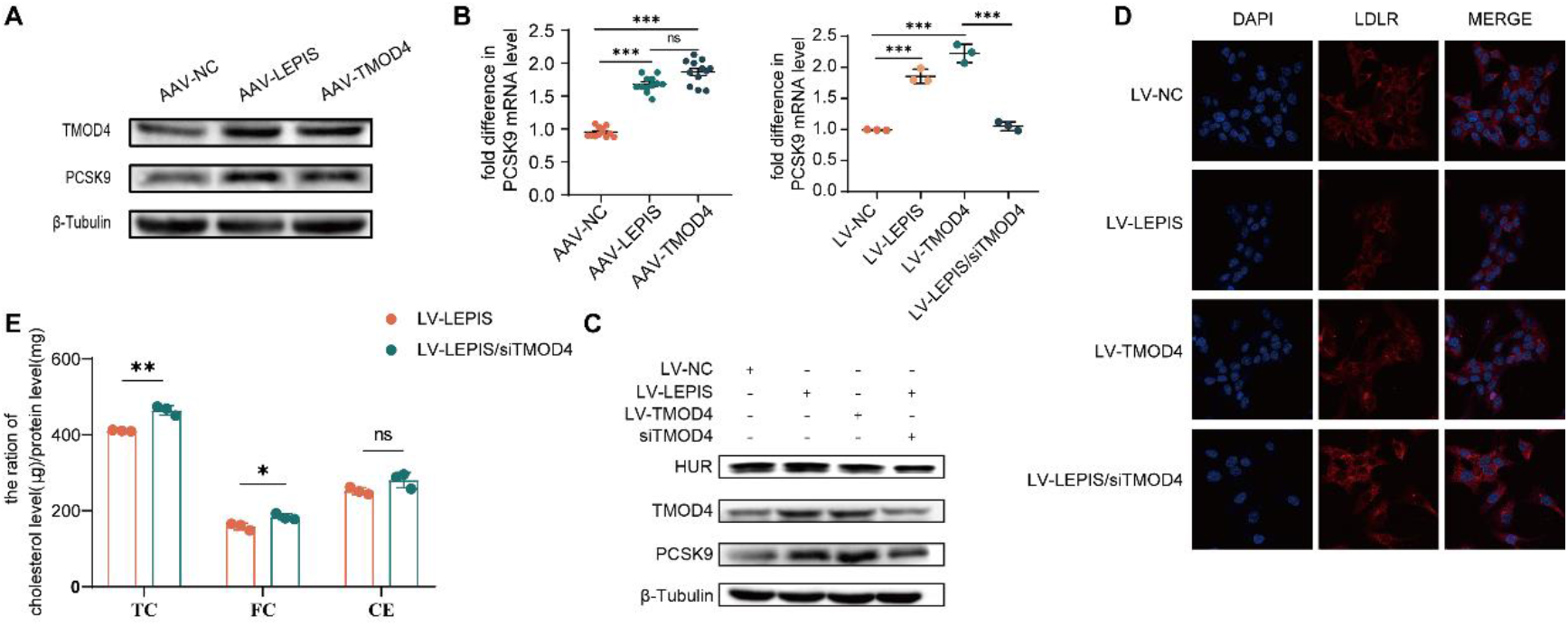
*LEPIS* increases *PCSK9* expression by enhancing *TMOD4* expression. (A) The protein levels of TMOD4 and PCSK9 in the livers of ApoE^-/-^ mice in each group; (B) Expression of *PCSK9* and *LDLR* in the HepG2 cells of LV-NC, LV-LEPIS, LV-TMOD4 and LV-LEPIS/siTMOD4; (C) The protein levels of TMOD4, PCSK9 and HuR in the HepG2 cells of each group; (D) Immunofluorescence for LDLR expression in HepG2 cells after overexpressing *LEPIS* and *TMOD4* (400×); (E) Levels of TC, FC and CE in HepG2 cells after the downregulation of *TMOD4*.

### *LEPIS* enhances *TMOD4* expression by facilitating the shuttling of HuR from the nucleus to the cytoplasm

LncRNAs regulate the transcription of downstream target genes by binding to RNA-binding proteins, and this is a major way that lncRNAs exert their biological effects(Ding, Yin et al., 2021). Through RBPDB prediction, lncLEPIS was found to have 4 binding sites with HuR (ELAV1) **(Supplementary Figure 5).** Therefore, we speculated that *LEPIS* regulates the expression of *TMOD4* through HuR. HuR is a multifunctional RNA-binding protein that is known to participate in various cellular processes, including cell signaling, RNA processing, and the regulation of gene expression(Ding et al., 2021, Eberhardt, Doller et al., 2012, Grammatikakis, Abdelmohsen et al., 2017). To investigate whether LEPIS could bind to HuR, RNA immunoprecipitation (RIP) was performed using an anti-HuR antibody, and any RNA bound to the HuR protein could thus be captured. RIP-qPCR analysis confirmed that there was a strong physical interaction between LEPIS and HuR (Figure 6A). And there was a weak interaction between LEPIS and TMOD4. which was abolished after *HuR* was silenced. To further verify whether *LEPIS* regulates the expression of TMOD4 through HuR, HepG2 cells (LV-LEPIS/siHuR) with overexpressed *LEPIS* and knocked down *HuR* were constructed. The results showed that after silencing *HuR,* the level of *TMOD4* expression in LV-LEPIS/siHuR cells was lower than that in LV-LEPIS cells (Figure 6B). Therefore, LEPIS regulates TMOD4 expression by binding to HuR. To clarify the binding sites of HuR and LEPIS, regions were predicted **(Supplementary Figure 6)**. Crosslinking immunoprecipitation followed by qPCR (CLIP-qPCR) was used to investigate whether HuR was mainly enriched at region 341–359 in the nucleotide sequence of LEPIS (Figure 6C). Based on the LEPIS (region 341-359) validated by CLIP-qPCR, some probes were designed for further validation using an RNA electrophoretic mobility shift assay (RNA-EMSA). The results showed that HuR could bind to the 341-359 region of LEPIS, and the above effect was abolished when the 341-359 nucleotide sequence of LEPIS was mutated (Figure 6D).

**Figure 6.**
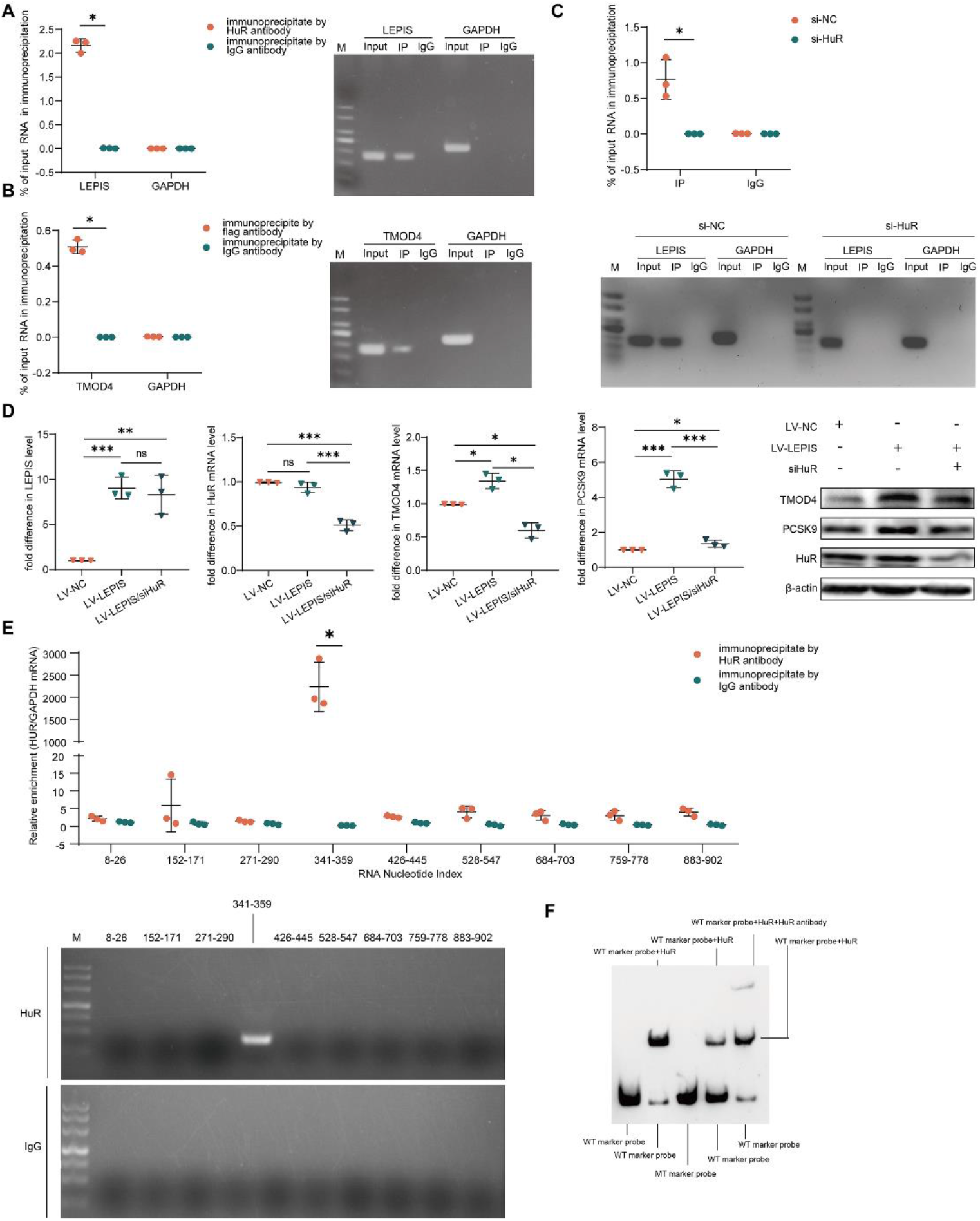
*LEPIS* enhances *TMOD4* expression by promoting the nucleocytoplasmic shuttling of HuR. (A) (Left) RT–PCR analysis of RNA immunoprecipitates by anti-HuR antibody. (Right) Representative images of the agarose gel of LEPIS RT–PCR products from input RNA samples, RNA immunoprecipitates by anti-HuR antibody, and RNA immunoprecipitates by IgG. (B) (Left) RT–PCR analysis of RNA immunoprecipitates by anti-flag antibody. (Right) Representative images of the agarose gel of LEPIS RT–PCR products from input RNA samples, RNA immunoprecipitates by anti-flag antibody, and RNA immunoprecipitates by IgG. (C) RT–PCR analysis of RNA immunoprecipitates by anti-flag antibody. Representative images of agarose gel of LEPIS RT–PCR products before and after the knockdown of HuR in HepG2 cells. (D) The expression of TMOD4, PCSK9 and HuR in LV-LEPIS HepG2 cells before and after *HuR* was silenced. (E) RT–PCR analysis of RNA immunoprecipitates (fragments of LEPIS) by anti-HuR antibody, and representative images of the agarose gel of fragments of the LEPIS RT–PCR products from input RNA samples, RNA immunoprecipitates by anti-HuR antibody, and RNA immunoprecipitates by IgG. (F) RNA-EMSA: The groups from left to right were wild-type labeled probe set, wild-type labeled probe + protein, mutant labeled probe + protein, wild-type labeled probe + wild competitive probe + protein, wild-type labeled probe +protein +antibody (Supershift).

However, surprisingly, we found that *LEPIS* overexpression did not increase the transcription and translation levels of HuR (Figure 5C). The idea that HuR increases the expression of target transcripts after shuttling from the nucleus to the cytoplasm and enhancing the stability of its target transcripts has been reported in the literature(Simion, Zhou et al.). Therefore, we further explored the effect of LEPIS on the cellular sublocalization of HuR and the way that HuR enhanced the expression of TMOD4. We examined the mRNA stability of TMOD4 under different intervention conditions. After actinomycin D intervention, the percentage remaining of *TMOD4* mRNA in LV-NC cells, LV-LEPIS cells and LV-LEPIS/siHuR cells was lower than before the intervention (Figure 7A). After actinomycin D intervention was performed for the same time or at the same concentration, the percentage remaining of TMOD4 mRNA in LV-LEPIS cells was higher than that in the LV-NC cells, while the percentage remaining of TMOD4 mRNA in the LV-LEPIS/siHuR cells was significantly lower than that in the LV-LEPIS cells (Figure 7A). This suggests that HuR, as an RNA-binding protein, binds to *TMOD4* mRNA and enhances its expression by promoting the stability of *TMOD4* mRNA. At the same time, we examined the intracellular distribution of HuR when LEPIS was overexpressed. Immunofluorescence and Western blot showed that compared with that in LV-NC cells, the expression of HuR was increased in the cytoplasm of LV-LEPIS cells and LV-LEPIS/siHuR cells (Figure 7B). This suggests that overexpression of LEPIS promotes the shuttling of HuR from the nucleus to the cytoplasm, and this process is independent of the total expression of HuR. To further explore whether HuR binds to *TMOD4* mRNA, RAP probes were designed for *TMOD4* mRNA **(Supplementary table 6)**, we performed an RNA antisense purification (RAP) experiment using antisense DNA tiling probes to pull-down *TMOD4* mRNA, followed by Western blot with anti-HuR antibody (Figure 7C-D). In addition, RNA immunoprecipitation was performed using HuR antibody, followed by detection of *TMOD4* mRNA (Figure 7E). RAP-Western Blot and RIP analysis confirmed that HuR interacts with *TMOD4* mRNA. Therefore, LEPIS enhances *TMOD4* mRNA stability by facilitating the shuttling of HuR from the nucleus to the cytoplasm and the interaction between HuR and *TMOD4* mRNA, thereby increasing TMOD4 expression.

**Figure 7.**
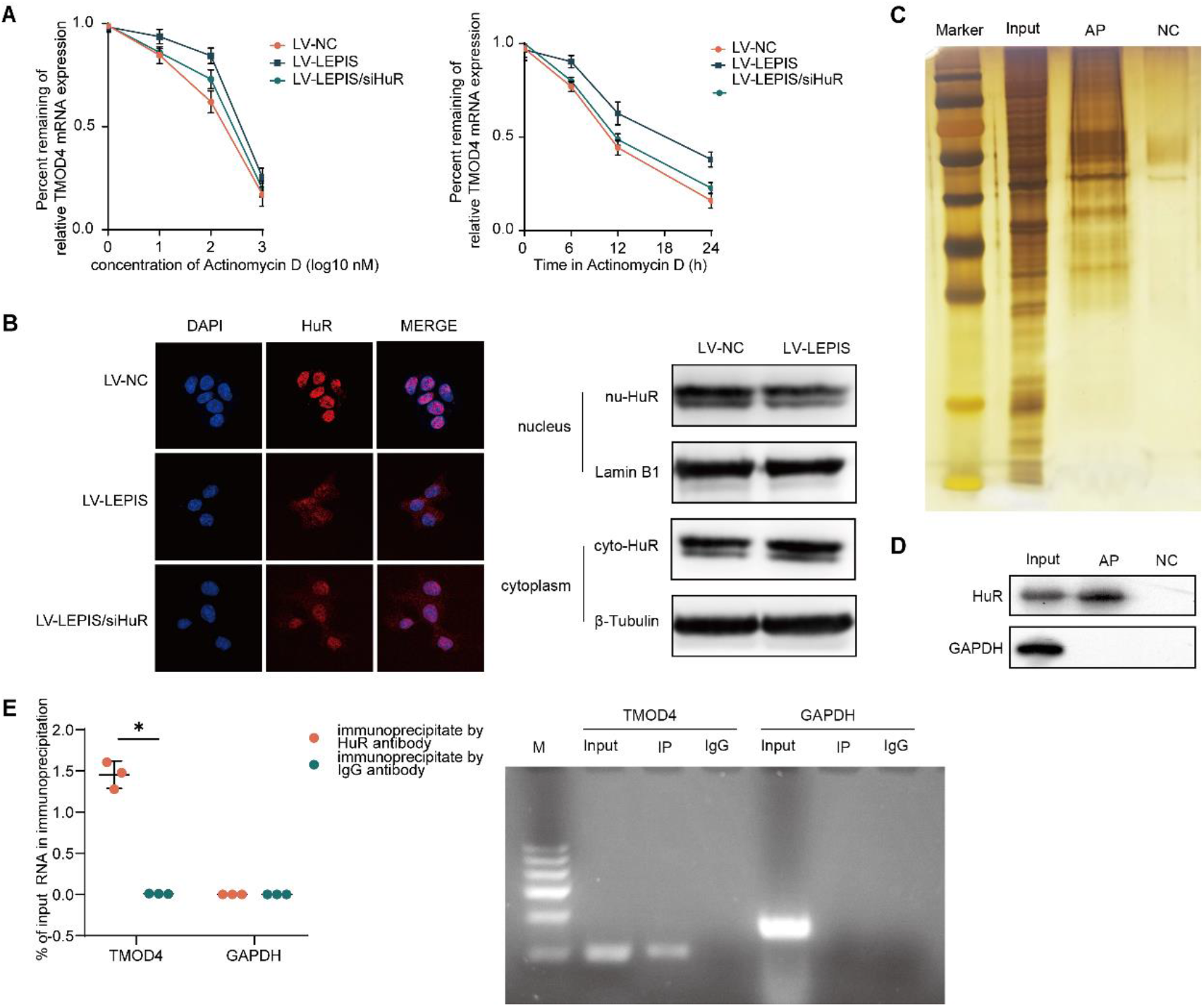
HuR binds to *TMOD4* mRNA and enhances its stability. (A) (Left) The percentage remaining of *TMOD4* mRNA in HepG2 cells treated with different concentrations of actinomycin D for 12 hours; (right) the percentage remaining of *TMOD4* mRNA in HepG2 cells treated with 10 nM actinomycin D for different times. (B) (Left) Subcellular localization of HuR (observed by immunofluorescence) in HepG2 cells before and after the overexpression of *LEPIS*. (Right) Protein levels of HuR in the nucleus and cytoplasm of HepG2 cells before and after the overexpression of *LEPIS*. (C) The RAP products were subjected to protein silver staining, and the results indicated that the RAP experiment had captured the protein, which could be used for subsequent Western Blot. (D) The HuR protein in the RAP products was detected by Western Blot, and the results showed that *TMOD4* mRNA was bound to the HuR. (E) RT–PCR analysis of RNA immunoprecipitates by anti-HuR antibody and representative images of the agarose gel of *TMOD4* mRNA RT–PCR products from input RNA samples, RNA immunoprecipitates by anti-HuR antibody, and RNA immunoprecipitates by IgG.

### TMOD4 regulates PCSK9 expression by affecting the transcriptional activity of the *PCSK9* promoter

Overexpressing TMOD4 resulted in an increase in *PCSK9* and a decrease in *LDLR* both in vivo and in vitro. A dual-luciferase reporter assay indicated that overexpression of TMOD4 enhanced the transcriptional activity of the *PCSK9* promoter and this effect was eliminated by mutating the *PCSK9* promoter (Figure 8A). To further explore how TMOD4 regulates the expression of PCSK9, we decomposed the sequence of the *PCSK9* promoter into 8 sequence fragments. ChIP–qPCR suggested that TMOD4 was bound to the *PCSK9* promoter at site 7 (Figure 8B). Based on the *PCSK9* promoter (site 7) validated by CHIP-qPCR, some probes were designed for further validation using a DNA electrophoretic mobility shift assay (DNA-EMSA). The results showed that TMOD4 could bind to site 7 of the *PCSK9* promoter, and when the fragment was mutated, the above effect disappeared (Figure 8C). This confirmed that TMOD4 regulated the expression of PCSK9 by affecting the transcriptional activity of the *PCSK9* promoter.

**Figure 8.**
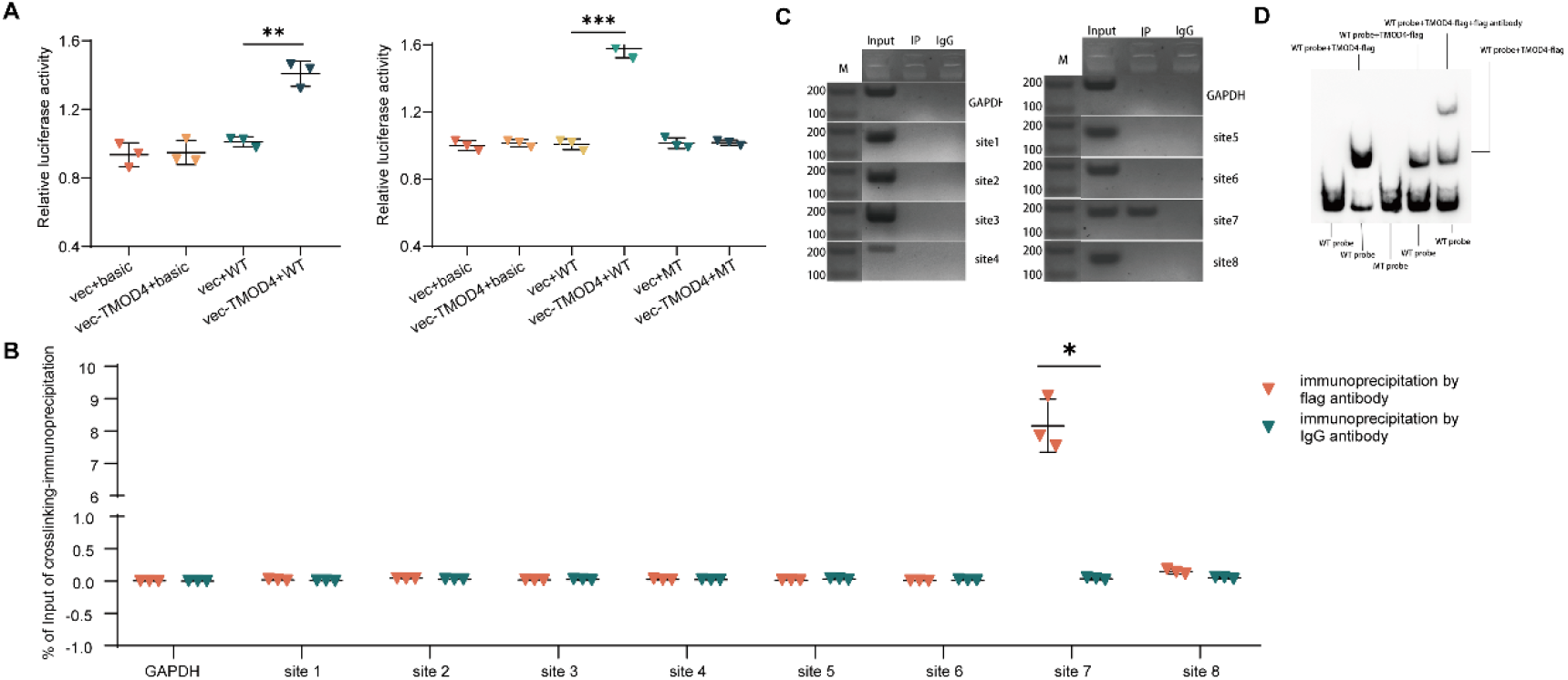
TMOD4 binds to the *PCSK9* promoter and affects its transcriptional activity. (A) Effect of TMOD4 overexpression on the transcriptional activity of the *PCSK9* promoter (luciferase reporter assay); (B) RT–PCR analysis of *PCSK9* promoter sequences of chromatin immunoprecipitation by anti-TMOD4 antibody; (C) Representative images of the agarose gel electrophoresis of PCR products; (D) DNA-EMSA: The groups from left to right were wild-type labeled probe, wild-type labeled probe + protein, mutant labeled probe + protein, wild-type labeled probe + wild competition probe + protein, wild-type labeled probe + protein + antibodies (Supershift).

## DISCUSSION

We mainly studied the uncharacterized lncRNA LEPIS, which plays an important role in AS. Our study confirmed that hepatic LEPIS increased the expression of PCSK9 and decreased the expression of LDLR by enhancing the expression of its target TMOD4; this in turn led to an impaired cholesterol uptake in hepatocytes and increased the level of LDL-C in plasma, which ultimately promoted the occurrence and progression of AS. The in vitro conclusions were consistent with those in the in vivo studies. We further found that LEPIS does not directly bind to its target TMOD4 but enhances the expression of *TMOD4* by interacting with the RNA-binding protein HuR and facilitating the shuttling of HuR from the nucleus to the cytoplasm. Overexpressing TMOD4 upregulates the transcriptional activity of *PCSK9* by binding to its promoter, ultimately leading to an impairment in hepatic cholesterol uptake. In addition, we also clearly elucidated the binding site of HuR to LEPIS. As a result, LEPIS is expected to become an effective new target for the prevention and treatment of AS, and the study also indicates that RNA-binding proteins may be closely related to AS **(Graphical abstract)**.

At present, there is enough evidence to confirm the key role of lncRNAs in atherosclerosis(Francesca, Di et al., 2019, Gao & Guo, 2021, Ym, Hy et al., 2021, Zhang, Yao et al., 2019). Notably, lncRNAs regulate the entire process of the formation and development of atherosclerosis. lncRNAs exert biological effects by regulating endothelial cells, macrophages, and vascular smooth muscle cells, which partly constitute the composition of plaques(Francesca et al., 2019). Moreover, some lncRNAs have even been regarded as novel diagnostic biomarkers for large atherosclerotic stroke(Zhang, Wang et al., 2021). In the past few decades, sufficient atherosclerosis-related lncRNAs have been discovered, and those related to cholesterol homeostasis include lincRNA-DYNLRB2-2, RP5833A20.1, HOXC-AS1, DYN-LRB2-2, MeXis, DAPK-IT1, and RP11-728F11.4(Huang, Hu et al., 2016, Kang, Hu et al., 2016, Yan-Wei, Jun-Yao et al., 2014, Zz, Sr et al., 2019). However, RBPs have not been well studied in atherosclerosis(Grammatikakis et al., 2017, Srikantan & Gorospe, 2012). A few RBPs, such as HuR, EWSR1, and KSRP, have been shown to affect atherosclerosis and the homeostasis of cholesterol(Ding et al., 2021). The interaction between lncRNAs and RBPs mainly occurs in three ways. First, RBPs regulate the expression of lncRNAs; second, lncRNAs regulate the expression of target genes by interacting with RBPs; and third, lncRNAs regulate the activity of their specific RBPs(Ding et al., 2021). In this study, *LEPIS* regulates the expression of the target gene *TMOD4* by interacting with HuR, so both LEPIS and HuR are important regulators of AS. In this process, the interaction of LEPIS with HuR did not involve changes in the levels of HuR transcription and translation, since the overexpression of LEPIS did not result in an enhancement or decrease in the level of HuR. Promoting the shuttling of HuR from the nucleus to the cytoplasm elevates the level of the target TMOD4, while a macrophage-specific lncRNA MAARS tethers HuR in the nucleus preventing its cytosolic shuttling(Simion et al.). Previous studies on HuR and AS have found that HuR is expressed in endothelial cells, macrophages, and smooth muscle cells, but its role in AS is highly dependent on the specificity of cell types(Meng et al., 2020, Srikantan & Gorospe, 2012). Our study is the first to investigate the expression of HuR in hepatocytes and its effect on AS, and the transport of HuR from the nucleus to the cytoplasm enhances the effect of *TMOD4* mRNA stability in a way similar to how HuR is regulated downstream in other cells and diseases.

Tropomodulin was originally discovered to be a protein that binds with tropomyosin to regulate the polymerization and depolymerization of actin(Weber, Pennise et al., 1994). Studies have suggested that the propionylation of TMOD3 is associated with platelet activation and arterial thrombosis(Xu, Jiang et al., 2020). The study pointed out that knocking out TMOD1 in systemic tissues other than the heart caused TG and HDL-C metabolism to be abnormal in mice, and this may be a cause of AS(Ding & Xu, 2015). In addition, TMOD1 in macrophages induces the formation of atherosclerotic plaques under certain conditions(Tursuntoheti, Sung et al., 2018). In an exon association study, *TMOD4* (rs115287176) was found to be significantly associated with myocardial infarction and hypertension(Yamada, Kato et al., 2018). Currently, the regulation of lipid metabolism and AS by tropomyosin has rarely been studied, and its mechanism is unclear. Both *LEPIS* and *TMOD4* are located on chromosome 1 and have overlapping segments, and this is a reason that the regulatory effect of *TMOD4* on cholesterol should be explored. Our study is the first to investigate the association of *TMOD4* in hepatocytes with cholesterol homeostasis. Our results suggest that *LEPIS* and *TMOD4* have the same effect on cholesterol homeostasis, as both lead to lower cellular cholesterol levels. Most of the previous studies on the correlation between tropomodulin and cholesterol metabolism were achieved through actin and focused on cardiomyocytes, macrophages, etc(Sedgwick, Balmert et al., 2018, Singh, Haka et al., 2019, Yamashiro, Gokhin et al., 2012). In contrast, this study found that TMOD4 binds to the *PCSK9* promoter region and regulates its transcriptional activity, thereby affecting LDLR recycling and plasma LDL-C levels.

Since cholesterol has many important functions in the physiological environment, disorders of cholesterol metabolism can lead to some congenital and acquired diseases, including familial hypercholesterolemia, Tangier disease, sitosterol disease and arteriosclerotic cardiovascular disease, which are all closely related to AS(Li, Yu et al., 2021). The homeostasis of cholesterol is inseparable from its synthesis, uptake, efflux and esterification(Luo et al., 2020). In this study, the increase in LDL-C in plasma was mainly due to impaired cholesterol uptake in hepatocytes, and the final targets of LEPIS and TMOD4 were PCSK9 and LDLR. In cholesterol homeostasis, LDLR primarily mediates the uptake of cholesterol-containing LDL particles from the blood by peripheral cells(Dandan, Han et al., 2021). The PCSK9-LDLR complex is directed to lysosomes and degraded, blocking LDLR recycling and the endocytosis of LDL particles(Luo et al., 2020). Indeed, further studies on NCP1L1-mediated cholesterol uptake in the gut lumen of mice would be helpful to thoroughly understand the effect of cholesterol uptake on plasma LDL-C levels. In addition, in this study, overexpressing *LEPIS* and *TMOD4* not only caused changes in PCSK9 and LDLR but also upregulated the expression of *ApoA2* and downregulated the expression of *ABCA1, ACAT2, ABCG5*, and *ABCG8*. These changes may also be related to cholesterol homeostasis and should be further studied.

In general, circulating HDL-C levels are inversely associated with the risk of cardiovascular disease(Sun, Guo et al., 2021). In this study, the levels of HDL-C in plasma were significantly elevated in AAV-LEPIS mice and AAV-TMOD4 mice, and this did not indicate that we challenged the notion that HDL-C has an antiatherosclerotic effect. Multiple current studies in mice and nonprimates have shown that PCSK9 is closely related to the metabolism of HDL-C(Choi, Aljakna et al., 2013a, Filippatos, Kei et al., 2018). Choi S et al. demonstrated that PCSK9 inactivation reduces the levels of HDL-C in serum and reduces the capacity of cholesterol efflux(Choi, Aljakna et al., 2013b). It has been reported that overexpressing *ApoA2* increases the levels of HDL-C levels animal models. The elevated overexpression of *PCSK9* and *ApoA2* in this study may also partly explain the elevated HDL-C in plasma(Bandarian, Daneshpour et al., 2016, Paththinige, Sirisena et al., 2017). Studies have found that a 30-70% increase in circulating HDL-C does not reduce recurring cardiovascular events in patients with ACS(Barter, Philip et al., 2007, Schwartz, Olsson et al., 2012, Sun et al., 2021). Madsen CM et al showed that both very high and very low HDL-C were associated with increased all-cause mortality(Neves, Batuca et al., 2021). In addition, we clearly know that HDL-C alone cannot accurately predict the risk of cardiovascular disease and that HDL-C dysfunction and remodeling are also very important.

Collectively, this study reveals an important regulatory role of the *LEPIS-HuR-TMOD4* axis in cholesterol homeostasis and AS. This not only deepens our understanding of the pathogenesis of AS but also once again emphasizes the importance of cholesterol homeostasis in AS. The study also enriches the related research on RBPs and AS. Both LEPIS and HuR are promising important targets for the prevention and treatment of AS, and more in-depth exploration is still necessary.

## MATERIALS AND METHODS

### Adeno-associated virus construction and effect verification

The LEPIS cDNA and TMOD4 cDNA were amplified by PCR and cloned into the pAV-CMV Globin Intron-MCS-P2A-EGFP-SV40polyA vector, and the correct sequences of the LEPIS gene and TMOD4 gene in this construct were verified by sequencing. This construct (referred to as AAV-LEPIS and AAV-TMOD4) and the vector (AAV-NC) were used to transfect the mice. The efficient overexpression of LEPIS and TMOD4 was verified by quantitative RT-PCR.

### Adeno-associated virus construction and effect verification

The LEPIS cDNA and TMOD4 cDNA were amplified by PCR and cloned into the pLVX-EF1a-Puro-WPRE-CMV-MCS vector, and the correct sequence of the LEPIS gene and TMOD4 gene in this construct was verified by sequencing. The construct pLVX-EF1a-Puro-WPRE-CMV-TMOD4-3FLAG was referred to as LV-TMOD4, and the construct pLVX-EF1a-Puro-WPRE-CMV-VPS72-205 was referred to as LV-LEPIS. LV-LEPIS, LV-TMOD4 and the control vector (referred to as LV-NC) were used to transfect cultured HepG2 cells. Efficient LEPIS and TMOD4 overexpression in HepG2 cells was verified by quantitative RT-PCR.

### Animals and their diet

The use of ApoE^-/-^ mice in this study was approved by the Animal Experiment Committee of Nanfang Hospital at Southern Medical University (Guangzhou, China). Forty-five ApoE^-/-^ mice aged 6-8 weeks were randomly divided into the following groups: the control group (AAV-NC), LEPIS overexpression group (AAV-LEPIS), and TMOD4 overexpression (AAV-TMOD4) group. The mice were fed a high-fat diet (HFD) for 8 weeks, and then three mice were randomly selected from each group and their aortas were extracted and stained with Oil Red O to determine AS. Subsequently, mice in each group were injected with different adeno-associated viruses (10^11^ pfu) through their tail veins. After continuing the HFD for 4 weeks, blood samples, liver tissue, and aorta of the mice were collected for the next step.

ApoE^-/-^ mice on a C57BL/6 background were obtained from Vital River Laboratory Animal Technology Co. Ltd. (Beijing, China), and the HFD (15% fat and 1.2% cholesterol) was purchased from Guangdong Provincial Animal Medicine Experimental Center (Guangzhou, China).

### Collection of blood samples, liver tissue and aorta

Mice were sacrificed 4 weeks after the tail vein injections, and the blood samples, liver tissue and aorta were quickly frozen in liquid nitrogen and stored in a −80°C freezer. Before the tissue was harvested, all the mice were weighed, and the heart and whole peripheral blood vessels of the mice were perfused.

### Lipid detection

Serum was collected to measure the levels of lipids, including triglycerides (TGs), total cholesterol (TCHO), LDL-C and HDL-C (Nanjing Jiancheng Bioengineering Institute, China).

### Oil Red O staining and imaging

To detect the burden of AS in mice, after the mice were sacrificed, the thoracic and abdominal cavities of the mice were opened, and then the section from the aortic root to the iliac artery was extracted and opened longitudinally. After staining with the ready-made Oil Red O staining solution (Yuan Ye, China, S19039), photos were taken with a digital camera against a blue background. Oil red O staining was performed on aortic root cross-sections and liver tissues, and photographs were taken using a Fluorescence Inversion Microscope System (Olympus CellSens Dimension, Japan).

### HE staining, Masson staining, EVG staining and immunofluorescence

When the mice were sacrificed, the aortic root tissue was collected, fixed with formalin and made into paraffin sections (5 μm) and frozen sections (5 μm). The above sections were subjected to HE staining (marking lipid core), Masson staining (marking collagen fibers), and EVG staining (marking elastic fibers). In addition, immunofluorescence tests with the aortic root tissue were performed using anti-CD68 antibody (Cell Signaling Technology, USA, 9778S) which labels macrophages and anti-aSMA antibody (Cell Signaling Technology, USA, 192455) which labels vascular smooth muscle cells. It was best to obtain images within 7 days of staining using a Japanese Olympus BX63 microscope.

### Cell culture and cell transfection

HepG2 cells (China Center for Type Culture Collection, China) were cultured using DMEM+10% fetal bovine serum in an incubator at 37°C and CO2 at 5%. Cells were transfected using Invitrogen Lipofectamine 2000 (Thermo Fisher Scientific, USA, 11668-019) according to the transfection manual. When transfecting lentivirus, the virus stock solution can be directly diluted with medium and then replaced. According to the different functions of the transfected lentivirus, HepG2 cells were divided into LV-NC, LV-LEPIS, and LV-TMOD4. The total RNA of cells was collected 48 hours after transfection or the total protein of cells was collected after 72 hours of transfection, and the next experiment was carried out.

### Immunofluorescence of cells

When the density of HepG2 cells in the confocal dish was 50-60%, immunofluorescence staining was performed using anti-LDLR antibody (Abcam, Cambridge, ab30532) and anti-HuR antibody (Cell Signaling Technology, USA, 12582S). The secondary antibody was purchased from Hangzhou Fude Biological Technology CO., LTD (China, FD0129). Images were acquired using a Leica LAS X (Leica, Germany).

### Cholesterol concentration determination

Cholesterol levels were measured in homogenized liver tissue and collected from treated HepG2 cells. Total cholesterol and total free cholesterol were detected using kits (Applygen, China, E1015-50 and E1016-50) according to the instructions, and the levels of cholesterol esters were obtained by calculations. The measured cholesterol concentrations were corrected for the concentration per mg protein.

### RNA isolation, reverse transcription, and quantitative real-time PCR

Total RNA was extracted from mouse liver tissue or treated HepG2 cells using Invitrogen TRIzol (Thermo Fisher Scientific, USA, 15596026). cDNA synthesis was performed in a 10 µl system using 5×Evo M-MLVRT Master Mix (Rui Zhen, China, AG11706). Next, real-time quantitative PCR was performed using 2×SYBR Green Pro Taq HS Premix (Rui Zhen, China, AG11701). Using ACTIN or GAPDH as a control primer (Sangon Biotech, China), the ΔΔCt method was used to analyze its relative expression. The primers used in this study are listed in the **Supplementary Table 2**.

### Western blot analysis

HepG2 cells or liver tissue were lysed with a mixture of RIPA and protease inhibitors (100:1). Protein concentrations were determined using the bicinchoninic acid (BCA) kit (Thermo, USA, 23227). After electrophoresis, electrotransport, blocking, and incubation with primary and secondary antibodies was performed, the protein was fully contacted with ECL luminescent solution (Hangzhou Fude Biological Technology CO., LTD, China, FD8000), and finally, the image of the target protein was obtained using the Kodak Image Station 2000MM imaging system. The primary antibodies included TMOD4 antibody (CUSABIO, China, CSB-PA609953ESR2HU); HuR antibody (Cell Signaling Technology, USA, 12582S); PCSK9 antibody (Cell Signaling Technology, USA, 85813S); β-tubulin antibody (Cell Signaling Technology, USA), 2146S); and GAPDH antibody (Cell Signaling Technology, USA, 5174S). The secondary antibody was purchased from Hangzhou Fude Biological Technology CO., LTD (China, FDG007).

### RNA immunoprecipitation

Nuclear extracts from HepG2 cells were prepared and incubated with HuR antibody (Cell Signaling Technology, USA, 12582S), anti-flag antibody (Cell Signaling Technology, USA, 14793S) or nonspecific mouse IgG (Cell Signaling Technology, USA, 3900S) overnight at 4°C. After adding Protein G Plus Protein A agarose beads (Calbiochem, Germany), the mixture was incubated at 4°C for 1 hour, the beads were collected using a magnetic stand, and the supernatant was discarded. After washing and elution, RNA was extracted using TRIzol (Takara, China), and RT–PCR was performed. The primers used are listed in the **Supplementary Table 3**.

### RNA antisense purification (RAP)

Cultured HepG2 cells were cross-linked in 1% formaldehyde for 10 minutes, followed by quenching in 0.125 M glycine for 5 minutes. The cells were then lysed, and DNA was removed by DNase I digestion. Thereafter, cell lysates were incubated with TMOD4 mRNA probes to pull-down TMOD4 mRNA, and streptavidin magnetic beads were used to capture TMOD4 mRNA-bound RNA and protein. The levels of HuR protein were detected using western blot and anti-HuR antibody (Cell Signaling Technology, USA, 12582S). The probes used are listed in the **Supplementary Table 6**.

### Chromatin immunoprecipitation

Harvested HepG2 cells were cross-linked with formaldehyde and then quenched with glycine. Chromatin immunoprecipitation was performed using the ChIP kit (Axl-Bio, China) and anti-TMOD4 antibody (CUSABIO, China, CSB-PA609953ESR2HU) according to the instructions. DNA immunoprecipitated by anti-TMOD4 antibody (CUSABIO, China, CSB-PA609953ESR2HU) was amplified using sequence primers for fragments of the PCSK9 promoter and GAPDH. The primers are listed in the **Supplementary Table 5**.

### Crosslinking-immunoprecipitation

HepG2 cells were harvested and cross-linked using a 365 nm UV lamp (dose of 0.15 J/cm^2^). Immunoprecipitation was performed using a PAR-CLIP kit (Axl-Bio, China) and anti-HuR antibody (Cell Signaling Technology, USA, 12582S) according to the instructions. RT–PCR and agarose gel electrophoresis were performed after RNA extraction with TRIzol (Takara, China). The primers are listed in the **Supplementary Table 4**.

### Luciferase reporter assay

The PCSK9 gene promoter sequence was cloned into the pGL4.10 vector containing the firefly luciferase reporter gene, and pGL4.10 constructs with a truncated PCSK9 gene promoter sequence were generated. Cultured HepG2 cells were transfected with either of these plasmids (together with a pRL-TK plasmid that contained the Renilla luciferase gene to serve as a reference for transfection efficiency), and cotransfection was performed with either a plasmid to overexpress TMOD4 or a mutated plasmid or the plasmid vector (control); then, firefly luciferase activity was measured with standardization against the Renilla luciferase activity.

### Electrophoretic mobility shift assay (EMSA)

According to the LEPIS region verified by CLIP-qPCR and regions of the PCSK9 promoter validated by luciferase experiments, an EMSA probe set was designed for EMSA verification. The wild-type probes, mutant probes and wild-competing probes were designed, wherein both the wild-type probes and mutant probes were labeled with biotin. Nucleoprotein extracts were electrophoresed and analyzed according to the principle that protein-probe complexes migrate slowly during gel electrophoresis. The probes used are listed in the **Supplementary Table 7-8**.

### Statistical Analysis

Oil red O staining, HE staining, Masson and EVG staining and immunofluorescence images were quantified using the ImageJ system. The burden of AS was expressed by the area of aortic lesions/area of the entire aorta (%) and the area of aortic root lesions/aortic root cross-sectional area (%). All data are expressed as the mean ± SEM. Statistical analysis of the data was performed using GraphPad 8.0.1, and the comparison between a certain overexpression group and the control group was performed using an independent sample t test. P<0.05 was considered significant.

## AUTHOR CONTRIBUTIONS

Designing research studies: Kai Guo.

Conducting experiments: Ping Lv, Hangyu Pan, Kexin Hu.

Acquiring data, analyzing data: Ping Lv, Qinxian Li, Rongzhan Lin.

Providing reagents: Kai Guo, Zhigang Guo, Shaoyi Zheng.

writing the manuscript: Ping Lv, Kai Guo.

## ACKNOWLEDGMENTS

This work was supported by the Youth Fund of National Natural Science Foundation of China (81900398), the National Natural Science Foundation of China (82170474), Guangdong Basic and Applied Basic Research Foundation (2019A1515010666), Clinical Research Startup Program of Southern Medical University by High Level Construction Funding of Guangdong provincial Department of Education (LC2016PY002), and Clinical Research Program of Nanfang Hospital, Southern Medical University (2018CR501). The sponsors had no role in the study design, data collection and analysis, decision to publish or the manuscript.

## REFERENCES

Baigent C, Blackwell L, Emberson J, Holland LE, Reith C, Bhala N, Peto R, Barnes EH, Keech A, Simes J (2010) Efficacy and safety of more intensive lowering of LDL cholesterol: a meta-analysis of data from 170 000 participants in 26 randomized trialCholesterol Treatment Trialists’ (CTT) Collaboration Lancet 201037616701681 10.1016/S0140-6736(10)61350-5298822421067804. Lancet 376: 1670–1681

Bandarian F, Daneshpour MS, Hedayati M, Naseri M, Azizi F (2016) Identification of Sequence Variation in the Apolipoprotein A2 Gene and Their Relationship with Serum High-Density Lipoprotein Cholesterol Levels. Iranian Biomedical Journal 20

Barter, Philip J, Caulfield, Mark, Eriksson, Mats, Grundy, Scott M, Kastelein, John JP (2007) Effects of Torcetrapib in Patients at High Risk for Coronary Events. New England Journal of Medicine

Choi S, Aljakna A, Srivastava U, Peterson BR, Deng B, Prat A, Korstanje R (2013a) Decreased APOE-containing HDL subfractions and cholesterol efflux capacity of serum in mice lacking Pcsk9. Lipids in Health and Disease 12

Choi S, Aljakna A, Srivastava U, Peterson BR, Korstanje R (2013b) Decreased APOE-containing HDL subfractions and cholesterol efflux capacity of serum in mice lacking Pcsk9. Lipids in Health and Disease 12: 112

Dandan M, Han J, Mann S, Kim R, Mohammed H, Nyangau E, Hellerstein M (2021) Turnover Rates of the Low-Density Lipoprotein Receptor and PCSK9: Added Dimension to the Cholesterol Homeostasis Model. Arteriosclerosis Thrombosis and Vascular Biology 41: 2866–2876

Ding F, Xu D (2015) Changes in blood rheology and blood lipids in ApoE and Tmod1 double knockout mice. Chinese Journal of Hemorheology: 5

Ding Y, Yin R, Zhang S, Xiao Q, Zhu X (2021) The Combined Regulation of Long Non-coding RNA and RNA-Binding Proteins in Atherosclerosis. Frontiers in Cardiovascular Medicine 8

Eberhardt W, Doller A, Pfeilschifter J (2012) Regulation of the mRNA-Binding Protein HuR by Posttranslational Modification: Spotlight on Phosphorylation. Current Protein & Peptide Science 13: -

Filippatos TD, Kei A, Rizos CV, Elisaf MS (2018) Effects of PCSK9 Inhibitors on Other than Low-Density Lipoprotein Cholesterol Lipid Variables. Journal of Cardiovascular Pharmacology and Therapeutics 23: 3-12

Francesca F, Di GK, Lars M, Johnson JL (2019) Non-coding RNAs in cardiovascular cell biology and atherosclerosis. Cardiovascular Research: 12

Gao H, Guo Z (2021) LncRNA XIST regulates atherosclerosis progression in ox-LDL-induced HUVECs. Open Medicine 16: 117–127

Grammatikakis I, Abdelmohsen K, Gorospe M (2017) Posttranslational control of HuR function. Wiley Interdisciplinary Reviews-Rna 8

Guo, Kai, Lu, Dan, Zhao, Jinzhen, Liu, Jichen, Luo, Tiantian (2018) PSRC1 overexpression attenuates atherosclerosis progression in apoE(-/-) mice by modulating cholesterol transportation and inflammation. JOURNAL OF MOLECULAR AND CELLULAR CARDIOLOGY 116: 69–80

Huang C, Hu YW, Zhao JJ, Ma X, Wang Q (2016) Long Noncoding RNA HOXC-AS1 Suppresses Ox-LDL-Induced Cholesterol Accumulation Through Promoting HOXC6 Expression in THP-1 Macrophages. Dna & Cell Biology 35

Hussain A, Ballantyne CM (2021) New Approaches for the Prevention and Treatment of Cardiovascular Disease: Focus on Lipoproteins and Inflammation. Annual Review of Medicine 72

Johns DG, Almonte Y, Bautmans A, Campeau L-C, Cancilla MT, Chapman J, Crevecoeur I, Guetschow ED, Kauh EA, Lai E, Lanning CL, Lee AY, Li L, Mitchel YB, Stoch SA, Van Dyck K, Vanhoutte FP, Volckaert B, Wolford DG, Wood HB et al. (2021) The Clinical Safety, Pharmacokinetics, and LDL-Cholesterol Lowering Efficacy of MK0616, an Oral PCSK9 Inhibitor. Circulation 144: E573-E573

Kang CM, Hu YW, Nie Y, Zhao JY, Li SF, Chu S, Li HX, Huang QS, Qiu YR (2016) Long non-coding RNA RP5-833A20.1 inhibits proliferation, metastasis and cell cycle progression by suppressing the expression of NFIA in U251 cells. Molecular Medicine Reports 14: 5288-5296

Karagiannis AD, Liu M, Toth PP, Zhao S, Agrawal DK, Libby P, Chatzizisis YS (2018) Pleiotropic Anti-atherosclerotic Effects of PCSK9 Inhibitors From Molecular Biology to Clinical Translation. Current Atherosclerosis Reports 20

Knowles JW, Howard WB, Karayan L, Baum SJ, Wilemon KA, Ballantyne CM, Myers KD (2017) Access to Nonstatin Lipid-Lowering Therapies in Patients at High Risk of Atherosclerotic Cardiovascular Disease. Circulation 135: 2204–2206

Li H, Yu XH, Ou X, Ouyang XP, Tang CK (2021) Hepatic cholesterol transport and its role in non-alcoholic fatty liver disease and atherosclerosis. Progress in Lipid Research 83: 101109

Luo J, Yang H, Song BL (2020) Mechanisms and regulation ofcholesterol homeostasis. Nature Reviews Molecular Cell Biology

Meng Q, Pu L, Luo X, Wang B, Liu B (2020) Regulatory Roles of Related Long Non-coding RNAs in the Process of Atherosclerosis. Frontiers in Physiology 11: 564604

Mengqiu, Wei, Peng, Li, Kai, Guo (2020) The impact of PSRC1 overexpression on gene and transcript expression profiling in the livers of ApoE ?/? mice fed a high-fat diet. Molecular and Cellular Biochemistry 465: 125–139

Neves M, Batuca JR, Alves JD (2021) The role of high-density lipoprotein in the regulation of the immune response: implications for atherosclerosis and autoimmunity. Immunology

Paththinige CS, Sirisena ND, Dissanayake V (2017) Genetic determinants of inherited susceptibility to hypercholesterolemia – a comprehensive literature review. Lipids in Health & Disease 16

Schwartz GG, Olsson AG, Abt M, Ballantyne CM, Tiroch K (2012) Effects of Dalcetrapib in Patients with a Recent Acute Coronary Syndrome. New England Journal of Medicine 367

Sedgwick A, Balmert MO, D’Souza-Schorey C (2018) The formation of giant plasma membrane vesicles enable new insights into the regulation of cholesterol efflux. Experimental Cell Research 365: 194–207

Simion V, Zhou H, Haemmig S, Pierce JB, Mendes S, Tesmenitsky Y, Pérez-Cremades D, Lee JF, Chen AF, Ronda N A macrophage-specific lncRNA regulates apoptosis and atherosclerosis by tethering HuR in the nucleus. Nature Communications

Singh RK, Haka AS, Bhardwaj P, Zha X, Maxfield FR (2019) Dynamic Actin Reorganization and Vav/Cdc42-Dependent Actin Polymerization Promote Macrophage Aggregated LDL (Low-Density Lipoprotein) Uptake and Catabolism. Arteriosclerosis Thrombosis and Vascular Biology 39: 137–149

Srikantan S, Gorospe M (2012) HuR function in disease. Frontiers in Bioscience 17: 189

Sun L, Guo M, Xu C, Qiao X, Hua Y, Tuerhongjiang G, Lou B, Li R, Bai X, Zhou J, Wu Y, She J, Yuan Z (2021) HDL-C/apoA-I Ratio Is Associated with the Severity of Coronary Artery Stenosis in Diabetic Patients with Acute Coronary Syndrome. Disease Markers 2021

Tang Y, Li S-L, Hu J-H, Sun K-J, Liu L-L, Xu D-Y (2020) Research progress on alternative non-classical mechanisms of PCSK9 in atherosclerosis in patients with and without diabetes. Cardiovascular Diabetology 19

Tokgozoglu L, Libby P (2022) The dawn of a new era of targeted lipid-lowering therapies. European heart journal

Townsend N, Kazakiewicz D, Wright FL, Timmis A, Vardas P (2022) Epidemiology of cardiovascular disease in Europe. Nature reviews Cardiology

Tursuntoheti A, Sung LA, Yao W (2018) Tmod1 of macrophages is involved in the regulation of atherosclerotic plaque formation induced by the combination of high cholesterol and high matrix stiffness. In National Biomechanics Academic Conference and National Biorheology Academic Conference in China

Wang J-K, Li Y, Zhao X-L, Liu Y-B, Tan J, Xing Y-Y, Adi D, Wang Y-T, Fu Z-Y, Ma Y-T, Liu S-M, Liu Y, Wang Y, Shi X-J, Lu X-Y, Song B-L, Luo J (2022) Ablation of Plasma Prekallikrein Decreases LDL Cholesterol by Stabilizing LDL Receptor and Protects against Atherosclerosis. Circulation

Weber A, Pennise CR, Babcock GG, Fowler VM (1994) Tropomodulin caps the pointed ends of actin filaments. The Journal of cell biology 127: 1627–35

Weber C, Noels H (2011) Atherosclerosis: current pathogenesis and therapeutic options. Nature Medicine 17: 1410–1422

Xu Y, Jiang H, Li L, Chen F, Liu J (2020) Branched-Chain Amino Acid Catabolism Promotes Thrombosis Risk by Enhancing Tropomodulin-3 Propionylation in Platelets. Circulation 142

Yamada Y, Kato K, Oguri M, Horibe H, Fujimaki T, Yasukochi Y, Takeuchi I, Sakuma J (2018) Identification of 13 novel susceptibility loci for early-onset myocardial infarction, hypertension, or chronic kidney disease. International Journal of Molecular Medicine 42

Yamashiro S, Gokhin DS, Kimura S, Nowak RB, Fowler VM (2012) Tropomodulins: Pointed-end capping proteins that regulate actin filament architecture in diverse cell types. Cytoskeleton 69: 337–370

Yan-Wei H, Jun-Yao Y, Xin M, Zhi-Ping C, Ya-Rong H, Jia-Yi Z, Shu-Fen L, Yu-Rong Q, Jing-Bo L, Yan-Chao W (2014) A lincRNA-DYNLRB2-2/GPR119/GLP-1R/ABCA1-dependent signal transduction pathway is essential for the regulation of cholesterol homeostasis. Journal of Lipid Research 55: 681–697

Ym A, Hy A, Fd B (2021) LncRNA CDKN2B-AS1 in atherosclerosis: Friend or foe? International Journal of Cardiology

Zhang L, Yao D, Shen H, Gong K, Cardiology DO (2019) Research Progress in Effect of lncRNA in Atherosclerosis Lesions. Medical Recapitulate

Zhang S, Wang J, Qu MJ, Wang K, Zhu XY (2021) Novel Insights into the Potential Diagnostic Value of Circulating Exosomal IncRNA-Related Networks in Large Artery Atherosclerotic Stroke. Frontiers in Molecular Biosciences 8

Zz A, Sr A, Hj A, Xd A, Peng ZA, Jian LB (2019) The lncRNA DAPK-IT1 regulates cholesterol metabolism and inflammatory response in macrophages and promotes atherogenesis. Biochemical and Biophysical Research Communications 516: 1234–1241

